# Postpartum estrogen withdrawal induces deficits in affective behaviors and increases ΔFosB in D1 and D2 neurons in the nucleus accumbens core in mice

**DOI:** 10.1101/2022.09.08.505352

**Authors:** William B. Foster, Katherine F. Beach, Paige F. Carson, Kagan C. Harris, Brandon L. Alonso, Leo T. Costa, Roy C. Simamora, Jaclyn E. Corbin, Keegan F. Hoag, Sophia I. Mercado, Anya G. Bernhard, Cary H. Leung, Eric J. Nestler, Laura E. Been

## Abstract

In placental mammals, estradiol levels are chronically elevated during pregnancy, but quickly drop to prepartum levels following birth. This may produce an “estrogen withdrawal” state that has been linked to changes in affective states in humans and rodents during the postpartum period. The neural mechanisms underlying these affective changes, however, are understudied. We used a hormone-simulated pseudopregnancy (HSP), a model of postpartum estrogen withdrawal, in adult female C57BL/6 mice to test the impact of postpartum estrogen withdrawal on several behavioral measures of anxiety and motivation. We found that estrogen withdrawal following HSP increased anxiety-like behavior in the elevated plus maze, but not in the open field or marble burying tests. Although hormone treatment during HSP consistently increased sucrose consumption, sucrose preference was generally not impacted by hormone treatment or subsequent estrogen withdrawal. In the social motivation test, estrogen withdrawal decreased the amount of time spent in proximity to a social stimulus animal. These behavioral changes were accompanied by changes in the expression of ΔFosB, a transcription factor correlated with stable long-term plasticity, in the nucleus accumbens (NAc). Specifically, estrogen-withdrawn females had higher ΔFosB expression in the nucleus accumbens core. Using transgenic reporter mice, we found that this increase in ΔFosB occurred in both D1- and D2-expressing cells in the NAc core. Together, these results suggest that postpartum estrogen withdrawal impacts anxiety and motivation and increases ΔFosB in the NAc core.

## Introduction

Epidemiological studies consistently show that mood and anxiety disorders are more prevalent during life stages when estrogen levels are in flux. Indeed, puberty, pregnancy, and menopause are all associated with an increased incidence of mood and anxiety disorders, suggesting that estrogen fluctuations may confer additional vulnerability to psychiatric disorders (Soares and Zitek 2008). Among these life stages, the peripartum period stands out as extraordinarily volatile, both in terms of the associated estrogen changes, and the increased incidence of mental health disorders. By the end of the third trimester, estradiol levels increase up to 1000-fold (Hendrick, Altshuler, and Suri 1998). However, after birth and the expulsion of the placenta, estradiol levels quickly drop to below pre-partum levels and remain suppressed until ovulation resumes (McNeilly 2001). It has been hypothesized that this steep decline in estradiol levels results in an “estrogen withdrawal state” that contributes to the etiology of postpartum psychiatric disturbances (Hendrick, Altshuler, and Suri 1998; Bloch, Daly, and Rubinow 2003; Douma et al. 2005). This idea is supported by clinical studies demonstrating that the symptoms of postpartum depression can be attenuated by estradiol treatment (Sichel et al. 1995), and that experimentally-induced estrogen withdrawal can cause depressive symptoms in people with a history of postpartum depression (Bloch 2000).

To better understand the mechanisms by which postpartum estrogen withdrawal contributes to the etiology of psychological disorders, researchers have turned to rodent models. In particular, several groups have employed a hormone-simulated pseudopregnancy (HSP) model to directly test the impact of estrogen withdrawal on the brain and behavior. Initially developed in female Long-Evans rats (Galea, Wide, and Barr 2001), this model has successfully been replicated (Green, Barr, and Galea 2009) and extended to Sprague-Dawley rats (Schiller et al. 2013; Stoffel and Craft 2004; Suda et al. 2008; Navarre, Laggart, and Craft 2010; Baka et al. 2017), ICR mice (Zhang et al. 2016; Yang et al. 2017), and Syrian hamsters (Hedges et al. 2021), suggesting that it is a highly reproducible way to assess the impact of postpartum estrogen withdrawal across multiple species. In this model, females are ovariectomized and given daily injections that approximate estradiol levels in early- to mid-pregnancy, late pregnancy, and the postpartum period. In particular, females in an “estrogen-withdrawn” condition, which models the abrupt postpartum drop in estradiol, can be compared to females in an “estrogen-sustained” condition, in which estradiol continues to be administered during the simulated postpartum period, or vehicle-treated females. Consistent with the clinical literature, estrogen withdrawal following HSP results in a behavioral phenotype that may reflect depressed mood (Galea, Wide, and Barr 2001; Green, Barr, and Galea 2009; Schiller et al. 2013; Stoffel and Craft 2004; Suda et al. 2008; Navarre, Laggart, and Craft 2010) and/or heightened anxiety (Suda et al. 2008; Zhang et al. 2016; Yang et al. 2017) in rodents. In many cases, these behavioral changes can be prevented by continued administration of estradiol during the postpartum period, consistent with the data from postpartum human subjects.

The nucleus accumbens (NAc) is a behaviorally-relevant site of neuroplasticity following postpartum estrogen withdrawal. Located in the ventral striatum, the NAc receives projections from dopaminergic neurons in the midbrain ventral tegmental area (VTA), and this mesolimbic pathway plays a well-established role in motivated behaviors (Salamone et al. 2015), including maternal behaviors (Seip and Morrell 2009; Numan et al. 2005; Afonso et al. 2009; Numan 2007). This pathway also plays an important role in modulating mood/affect, as altered activity of NAc-projecting VTA neurons is associated with susceptibility to depression and anxiety behaviors in humans and animals (Russo and Nestler 2013; Tye et al. 2013, Lowes et al., 2021). Structural plasticity in the NAc has been linked to behavioral changes that reflect increased stress susceptibility, modeling a core symptom of mood and affective disorders (Christoffel et al. 2011; Bessa et al. 2013). Finally, the NAc is sensitive to estrogens (Creutz and Kritzer 2002; Mitra et al. 2003; Almey, Milner, and Brake 2015), making it a logical candidate for peripartum hormonal fluctuations to negatively impact the brain and behavior.

Within the NAc, the transcription factor ΔFosB is central to the molecular pathway that mediates structural and behavioral changes following chronic activation of the mesolimbic pathway (Nestler 2008), such as might be predicted following sustained hormonal changes during pregnancy and the postpartum period. Indeed, previous microarray data from postpartum hsd:ICR outbred mice shows increased *FosB* gene expression in the NAc (Zhao et al. 2014), and mutant mice lacking the *FosB* gene show deficits in parental and affective behaviors (Brown et al. 1996; Kuroda et al. 2008). However, the relationship between peripartum estrogen fluctuations and ΔFosB expression in the NAc is unknown. The present study therefore investigated the effect of estrogen withdrawal following HSP on affective and motivated behaviors, as well as the expression of ΔFosB, in C57BL/6 mice. We found that estrogen withdrawal following HSP increased anxiety-like behavior and decreased social motivation, while concurrently increasing ΔFosB in the NAc core. ΔFosB can have opposing effects on stress susceptibility in D1 vs. D2 neurons (Hamilton et al. 2018). Therefore, we also used transgenic fluorescent reporter mice to determine whether this increase in ΔFosB occurred selectively in NAc medium spiny neurons containing dopamine receptor 1 (D1) or dopamine receptor 2 (D2), and found that it occurred in both cell subtypes. Together, these findings suggest that peripartum estrogen fluctuations increase ΔFosB in D1 and D2 neurons in the NAc core, and that these changes are concurrent with increased anxiety and decreased social motivation during estrogen withdrawal.

## Methods

### Experimental Design and Hormone-Simulated Pseudopregnancy

A hormone-simulated pseudopregnancy (HSP) was used to test the impact of postpartum estrogen withdrawal on the behavior and the brain (**Figure 1**). Following recovery from ovariectomy (see below), subjects began the HSP injection regimen. On days 1-16, mice received daily subcutaneous injections containing a low dose (0.5 μg) of estradiol benzoate (Sigma) and a high dose (0.8 mg) of progesterone (Sigma) dissolved in 0.1mL cottonseed oil (Sigma). On days 17-23, mice received daily subcutaneous injections containing a high dose of estradiol benzoate (10 μg) dissolved in 0.1mL cottonseed oil (Sigma). On days 24-28, mice were divided into two experimental conditions: estrogen-withdrawn (“withdrawn”) females received daily vehicle injections, modeling the dramatic drop in estrogen that occurs during the postpartum period, while females in the estrogen-sustained (“sustained”) group continued to receive daily injections of a high dose (10 μg) of estradiol. In some experiments, these experimental groups were also compared to a vehicle control group that received oil injections throughout the duration of the HSP (“no hormone”). The doses and time course of the HSP hormone injections were initially chosen for their ability to elicit maternal responses in nulliparous, ovariectomized female rats (Galea, Wide, and Barr 2001), then modified for use in mice (Zhang et al. 2016). In rats, these doses produce serum concentrations that are supraphysiological for the high dose of estradiol, but are consistent with the level of progesterone reported during pregnancy (Bridges 1984; Garland et al. 1987; Galea, Wide, and Barr 2001).

**Figure 1:**
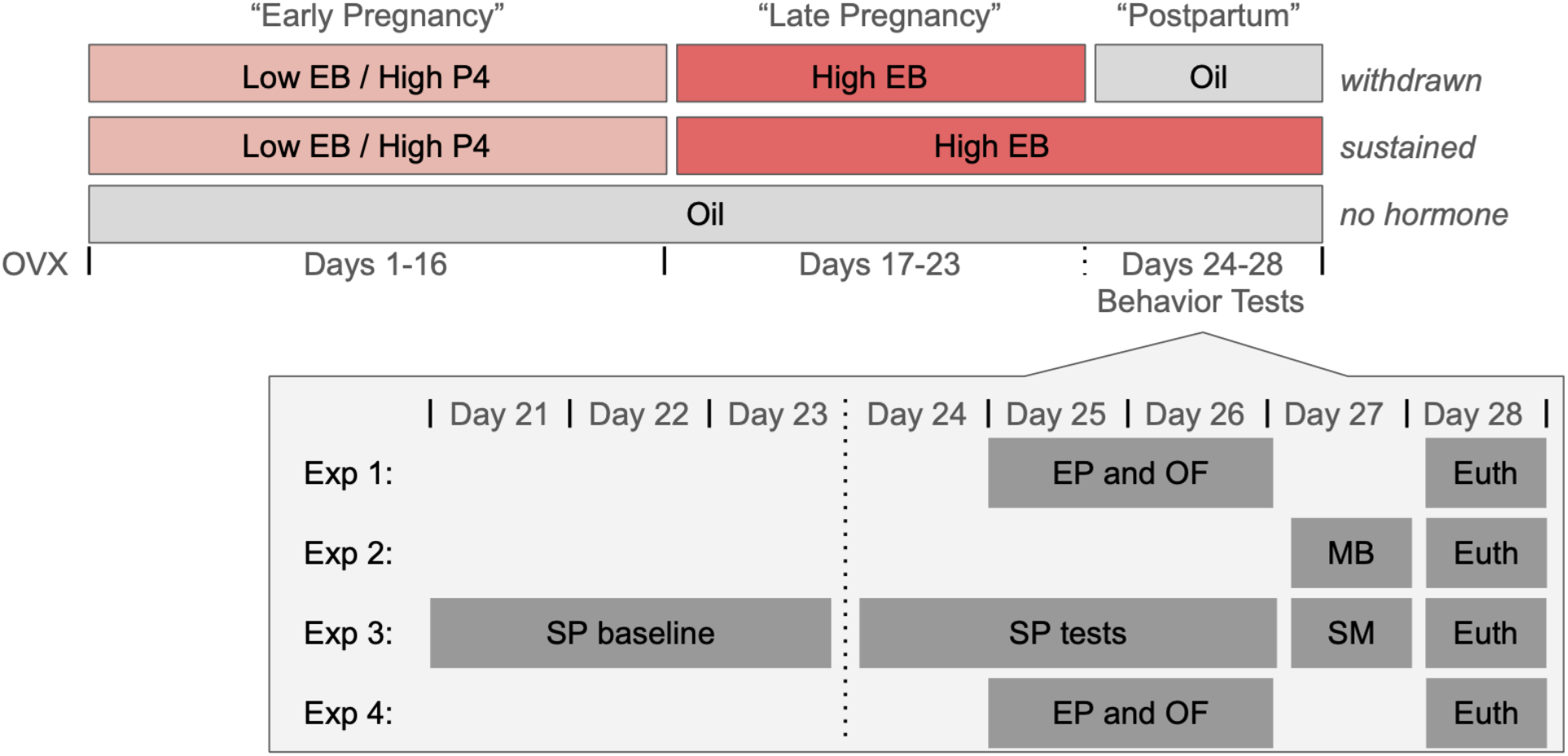
Hormone Simulated Pregnancy and Experimental Timeline. Following recovery from ovariectomy (OVX), subjects were assigned to one of three hormone simulated pregnancy conditions. Subjects in both the estrogen-withdrawn and estrogen-sustained conditions received daily injections containing a low dose of 17-β-estradiol (EB) and a high dose of progesterone (P4) on days 1-16, followed by injections containing a high dose of EB on days 17-23. On days 24-28, subjects in the withdrawn condition received oil injections, simulating the rapid drop in estradiol levels during the postpartum period. In contrast, subjects in the sustained condition continued to receive injections containing a high dose of EB throughout the simulated postpartum period. In some experiments, withdrawn and sustained subjects were compared to ovariectomized no hormone animals that received oil injections throughout the duration of the simulated pregnancy and postpartum period. On days 24-27, subjects across four experiments (Exp 1-4) underwent behavior testing in the elevated plus (EP), open field (OF), marble burying (MB), sucrose preference (SP), and social motivation (SM) tests. On day 28, subjects were euthanized (euth) and their brains were removed for histological processing.

The role of estrogen withdrawal following HSP on anxiety-like behaviors was assessed using elevated plus, open field, and marble burying tasks. Elevated plus and open field were assessed in the same cohort of subjects (Experiment 1), whereas marble burying was assessed in a separate cohort of subjects (Experiment 2), which only included estrogen-sustained vs. estrogen-withdrawn animals. The role of estrogen withdrawal following HSP on motivated behaviors was assessed using sucrose preference and social motivation tasks. Motivated behaviors were assessed in a separate cohort (Experiment 3) from either anxiety behavior cohort.

Following HSP (Day 28, 5 days post-withdrawal), subjects were euthanized and their brains were processed for FosB immunoreactivity (FosB-ir) in the NAc (core and shell). In a subset of animals, ΔFosB protein levels were measured in the NAc using Western blots. The results of these experiments indicated that estrogen withdrawal following HSP increased ΔFosB in the NAcC. However, ΔFosB may be expressed differentially, and have different functional properties, in D1 vs. D2 neurons in the NAc (Gagnon et al., 2017; Grueter et al., 2013). Therefore, a separate cohort of animals (Experiment 4) was used to test the role of estrogen withdrawal on ΔFosB expression following HSP in two different lines of female transgenic reporter mice on a C57BL/6 background. The use of these two mouse lines allowed for cell-type-specific localization of ΔFosB within functionally-distinct populations of D1 and D2 neurons in the NAc. Anxiety-like behavior in the elevated plus maze and open field tests was also assessed in these transgenic mice. These experiments only compared estrogen-sustained vs. estrogen-withdrawn animals.

### Subjects

A total of 90 mice were used in this study. Experiments 1 (*n* = 24), 2 (*n* = 16), and 3 (*n* = 24) used adult female C57BL/6 mice ordered at 8-12 weeks of age from Charles Rivers Laboratories (Wilmington, MA, USA). Experiment 4 used two different lines of adult female bacterial artificial chromosome (BAC) transgenic reporter mice on a C57BL/6 background. The first mouse line, *D1-tdtomato* (n = 8), expresses a red fluorescent protein under control of the D1 receptor promoter. The second line, *D2-GFP* (n = 8), expresses a green fluorescent protein under control of the D2 receptor promoter. Transgenic mice were shipped from Icahn School of Medicine at Mount Sinai to Haverford College at 7-13 weeks of age.

Stimulus mice (*n* = 10) for the social motivation test were juvenile female C57BL/6 mice, ordered at three weeks of age from Charles Rivers Laboratories. Prior to social motivation tests, juvenile mice were acclimated to being placed into a metal mesh cup within one of the holding chambers for five minutes for three consecutive days.

In all experiments, mice were housed in pairs in plexiglass cages (28cm x 17cm x 12cm) with aspen bedding, cotton nesting material, and unlimited access to food and water. All mice, including stimulus mice, were acclimated to being handled prior to the start of the experiment. Animal holding rooms were temperature- and humidity-controlled and maintained on a 12h/12h reversed light-dark cycle. All animal procedures were carried out in accordance with the National Institutes of Health Guide for the Care and Use of Laboratory Animals and approved by the Haverford College Institutional Animal Care and Use Committee.

### Ovariectomy

All surgery was conducted using aseptic surgical technique and under isoflurane anesthesia (2-3% vaporized in oxygen, Piramal, Bethlehem, PA, USA). Analgesic (Meloxicam, 5 mg/kg, Portland, ME, USA) was administered subcutaneously prior to the start of surgery and for three days postoperatively. For ovariectomies, subjects’ bilateral flanks were shaved and cleaned with three alternating scrubs of 70% ethanol and Betadine before being transferred to a temperature-regulated, sterile surgical field. Isoflurane anesthesia was maintained via a nose cone. Subjects’ ovaries were extracted via bilateral incisions of the smooth muscle under the flanks and then removed via cauterization of the uterine horn and blood vessels. Polydioxanone absorbable suture (Ethicon, Sommerville, NJ, USA) and wound clips (Fine Science Tools, Foster City, CA, USA) were used to close the smooth muscle and skin incisions, respectively.

### Behavior Testing

#### Elevated Plus

On day 25 or 26 (“postpartum” days 2 and 3), subjects in experiment 1 or 4 were tested for thigmotaxis behavior as an assay of anxiety using the elevated plus. The apparatus consists of a plus-shaped maze elevated 73 cm above the floor, with two open arms (51 × 11.5 cm) and two enclosed arms (51 × 11.5 × 39.5 cm). Mice were placed in the center of the maze facing a closed arm and allowed to freely explore the maze for five min under white light. The testing chamber was cleaned thoroughly with 70% EtOH between subjects. Tests were video recorded and the amount of time spent in each arm, as well as the total distance traveled and average velocity were quantified using Noldus Ethovision XT (version 14, Noldus Information Technology, Wageningen, The Netherlands).

#### Open Field

On day 25 or 26 (“postpartum” days 2 and 3), subjects in experiment 1 or 4 were tested for thigmotaxis behavior as an assay of anxiety using the open field. The apparatus consists of an open field (40.5 × 40.5 × 30cm) composed entirely of solid, light gray plastic (Maze Engineers, Glenview, IL, USA). Mice were placed in the center of the apparatus and allowed to freely explore for five min under white light. The testing chamber was cleaned thoroughly with 70% EtOH between subjects. Tests were video recorded and the amount of time spent in the center of the field, periphery of the field, as well as the total distance traveled and average velocity were quantified using Noldus Ethovision XT.

#### Marble Burying

On day 27 (“postpartum” day 4), subjects in Experiment 2 were tested for compulsive-like behavior using a marble burying test. Clean mouse cages were prepared with aspen bedding (approximately 5 cm deep) and 15 multicolor glass marbles (16-mm diameter) that had been cleaned with 70% EtOH. Marbles were evenly-distributed on the surface of the bedding, such that they were 2.5 cm from the sides of the cage and with ∼4 cm between each other. Prior to the initiation of HSP, subjects were tested for baseline marble burying behavior so that marble burying could be balanced across hormone conditions. On day 27 (“postpartum” day 4), subjects were again tested for their marble burying behavior. During both baseline tests and experimental tests, mice were placed into the prepared cage for 30 minutes and allowed to freely interact with the marbles. After 30 minutes, mice were carefully removed from the cage and the number of buried marbles was counted by two experimenters blind to the condition of the subjects. A marble was considered buried if more than two thirds of the marble was covered in bedding.

#### Sucrose Preference

Subjects in Experiment 3 were tested for motivation for palatable solutions, or possible anhedonia, using a sucrose preference test. On Days 21-23 (“late pregnancy”), all mice were acclimated to the two-bottle choice task. On these days, two bottles were available in each home cage: one bottle was filled with tap water and the other with 2% sucrose solution. Prior to testing, subjects were water deprived for 5 hr. Subsequently, each subject was individually housed in a clean testing cage with one bottle of tap water and one bottle of 2% sucrose solution for 3 hr before being returned to their home cage. Bottle side was carefully switched at the 1.5-hour mark to account for potential side bias. Each bottle was weighed before and after the 3-hr period on each day in order to determine fluid intake. On Days 24-26 (“postpartum” days 1-3), the 5-hr water deprivation followed by 3-hr two-bottle test procedure was repeated. Sucrose preference was calculated as sucrose solution consumption + water consumption / sucrose solution consumption x 100%.

#### Social Motivation

On Day 27 (“postpartum” day 4), subjects in Experiment 3 were tested for their motivation to investigate a social stimulus. The apparatus consists of two lateral chambers (21 × 19 × 29.5 cm) composed of solid, light gray plastic, and a clear plastic holding chamber (21 × 6.5 × 29.5 cm) and removable doors (Maze Engineers). Each of the lateral chambers held an inverted mesh cup (9.5 × 9.5 × 12 cm): one cup contained a toy black mouse toy (Play Fur Mice Cat Toy, Penn-Plax) and the other contained a juvenile (30 day old) female stimulus mouse, both of which were previously unfamiliar to the subjects. Subjects were placed in the holding chamber for 5 minutes of acclimation, after which the doors were lifted and the subject was allowed to freely explore the two stimulus chambers for 10 minutes before being returned to their home cage. To control for side bias, the placement of the social and non-social stimuli was alternated between subjects. The testing chamber was cleaned thoroughly with 70% EtOH between subjects. Tests were video recorded and the amount of time spent in each chamber, and the time spent interacting with either stimulus, and the amount of time spent facing the social stimulus was quantified using Noldus Ethovision XT. Interactions were defined as a mouse having its nose within 2 cm of stimulus containers while having its nose pointed in the direction of the container. Relative duration in the social floor was calculated as the time in floor of social stimulus / total time in each floor. Relative time in proximity to social stimulus was calculated as the time spent within 2 cm of social stimulus / total time in proximity to both stimuli. Relative time facing the social stimulus was calculated as the time oriented towards the social stimulus / total time oriented to both stimuli.

### Histology

#### Perfusion

On Day 27 (“postpartum” day 5), subjects were given an overdose of sodium pentobarbital (Euthanasia III, Patterson Vet Supply, Devens, MA, USA) and transcardially perfused with approximately 25 mL of cold 25 mM PBS (pH 7.4), followed by 50 mL of cold 4% paraformaldehyde (Electron Microscopy Sciences, Hatfield, PA, USA). Brains were immediately removed and post-fixed in 4% paraformaldehyde overnight and then cryoprotected for at least 48 hr in 30% sucrose in PBS. Coronal sections (35 to 40-μm) of brain tissue were sectioned on a cryostat (−20°C), collected in a 1:4 series, and stored at -20°C in cryoprotectant until immunohistochemical processing.

#### Immunohistochemistry and brightfield microscopy

Tissue from experiments 1 and 2 were processed for immunohistochemical detection of FosB proteins (ΔFosB and full-length FosB) in the NAc. Briefly, sections containing the NAc were removed from cryoprotectant and rinsed 5 × 5 min in 25 mM PBS. To reduce endogenous peroxidase activity, tissue sections were incubated in 0.3% hydrogen peroxide for 15 min. After 5 × 5 min rinses in PBS, sections were incubated in a monoclonal primary antibody against FosB (1: 2,000, (5G4) Rabbit mAb #2251, Cell Signaling) in 0.4% Triton-X 100 for 24 hr at room temperature. After incubation in primary antibody, sections were rinsed in five times in PBS and then incubated for 1 hr in biotinylated secondary antibody (anti rabbit, 1:600 Jackson Immunoresearch, West Grove, PA, USA) in PBS with 0.4% Triton-X 100. Sections were again rinsed five times in PBS and then incubated for 1 hr in avidin-biotin complex (4.5 μL each of A and B reagents/ml PBS with 0.4% Triton-X 100, ABC Elite Kit, Vector Laboratories, Burlingame, CA, USA). After rinsing in PBS, sections were incubated in 3,3’-diaminobenzidine HCl (0.2 mg/mL, Sigma) with hydrogen peroxide (0.83 μL/mL, Sigma) for 10 min, yielding an orange-brown product. The reaction was stopped by rinsing sections in PBS. Stained tissue sections were mounted onto subbed glass slides and allowed to air-dry overnight. Slides were then dehydrated in alcohols, cleared in xylenes, and cover slipped using Permount (Fisher Scientific, Waltham, MA, USA).

A light microscope (Nikon Eclipse E200) using a color camera (Diagnostic Instruments, Sterling Heights, MI, USA) and Spot Basic software was used to acquire images. 10X images of the bilateral NAc Shell and Core (2 unilateral images per animal) were acquired. Selection of sections for analysis was based on neuroanatomical matching to stereotaxic atlas plates containing the NAc (Bregma +1.10 mm, Paxinos and Franklin, 2019). Counting domains for the NAcc and NAcSh were consistently applied onto images using the anterior commissure as a landmark (see Figure 4). All cell counts were done manually by experimenters blind to the condition of the subject, and inter-rater reliability was confirmed. FosB-ir cell counts were also attempted in the medial preoptic area (MPOA), but FosB-ir was sparse in this brain region.

#### Double immunofluorescence and confocal microscopy

Tissue sections from Experiment 4 were processed for FosB immunofluorescence in either D1- or D2-expressing neurons in the NAc. Briefly, sections containing the NAc were removed from cryoprotectant and rinsed 5 × 5 min in 25 mM PBS. Sections were incubated in the appropriate primary antibodies (see below) in 0.1 PBTX for 24 hours at room temperature. After incubation in primary antibody, sections were rinsed five times in PBS and then incubated in the appropriate fluorescent secondary antibodies (see below) in 0.1% PBTX for one hour at room temperature in the dark. In tissue from *D1-tdtomato mice*, primary antibodies were Rabbit anti-FosB (1:1000, Abcam #184938) and Goat anti-RFP (1:750, Rockland #200-101-397) and secondary antibodies were Alexa 488 anti-Rabbit and Alexa 594 anti-Goat (1:500, Jackson Immunoresearch, #111-545-144 and #705-585-147). In tissue from *D2-GFP* mice, primary antibodies were Rabbit anti-FosB (1:1000, Abcam #184938) and Goat anti-GFP (1:5000, Abcam #5450) for the primary antibodies and secondary antibodies were Alexa 594 anti-Rabbit and Alexa 488 anti-Goat (1:500, Jackson Immunoresearch, #711-585-152 and #705-546-147). After five final rinses in PBS, tissue was mounted onto subbed glass slides and coverslipped using ProLong Gold Antifade Mountant (Invitrogen, Waltham, MA, USA). Slides were stored in the dark until imaging.

Double-immunofluorescent tissue was imaged using a confocal microscope (Nikon C1 Confocal Microscope) using EZ-C1 software (version 3.90, Nikon). 20X images of the bilateral NAc (2 unilateral images per animal) were acquired. Selection of sections for analysis was based on neuroanatomical matching to stereotaxic atlas plates containing the NAc (Bregma +1.10 mm, Paxinos and Franklin 2019). Consistent counting domains for the NAcC and NAcSh were consistently applied using the anterior commissure as a landmark (See **Figure 4**). Double-labeled cells were identified as having a FosB nuclear stain (green in *D1-tdtomato* animals and red in *D2-GFP* animals) co-localized with cytoplasmic staining for transgenic expression of either *tdtomato* (red) or *GFP* (green). This colocalization typically appeared as yellow (**Figure 5**). In tissue from *D1-tdtomato* mice, photomicrographs were post-processed using the “Find Edges” function in FIJI to more clearly differentiate D1 cells from background. All cell counts were done manually by at least two experimenters blind to the condition of the subject, and inter-rater reliability was confirmed.

#### Western Blots

A subset of estrogen-withdrawn (*n* = 6) and estrogen sustained (*n* = 6) animals in experiment 1 were given an overdose of Euthanasia III (Patterson Vet Supply) and sacrificed by rapid decapitation. 1-mm thick coronal sections containing the NAc were taken and bilateral tissue punches (1-mm diameter) were immediately collected from both brain areas. NAc punches included both the shell and core, but were biased towards the core (i.e., included the anterior commissure). Bilateral punches were flash-frozen and stored at -80°C until Western Blotting.

Bilateral tissue punches were homogenized in 1% sodium dodecyl sulfate (SDS) processing buffer and protein was quantified using the Bio-Rad protein DC assay (Bio-Rad Laboratories, Berkeley, CA). For each sample, 40 mg of total protein was loaded onto a 12-15% polyacrylamide gradient gel (Mini-PROTEAN TGX Precast Mini Gel; Bio-Rad Laboratories) and transferred to a nitrocellulose membrane following separation. Samples from each condition were counterbalanced across gels, and each sample was duplicated for technical replication. Membranes were blocked in 5% nonfat dried milk in TBS, incubated overnight in primary antibodies against FosB (1:1,000, Cell Signaling, Beverly MA) or Glyceraldehyde 3-phosphate dehydrogenase (GAPDH, 1:30,000, Millipore, Burlington, MA) diluted in Tris-buffered saline (TBS) + 0.1% Tween 20 (Sigma-Aldrich, St. Louis, MO) for 24 hr at room temperature. The following day, membranes were incubated in the appropriate horseradish peroxidase (HRP)-conjugated secondary antibodies (FosB anti-rabbit HRP, GAPDH anti-mouse HRP, Cell signaling) at 1:500 for 1 hr at room temperature. These primary antibodies have been used previously and produced bands at the appropriate molecular weights (Acaba et al. 2019). Following primary antibody incubation, membranes were developed for chemiluminescent detection using SuperSignal West Femto Maximum Substrate Sensitivity (Thermo Fisher Scientific, Waltham, MA). Chemiluminescence was visualized and quantified by an experimenter blind to the condition of the sample using FluorChem Software (Protein Simple, San Jose, CA). ΔFosB levels were normalized to GAPDH levels to control for equal loading of samples across wells.

### Statistical Analysis

Statistics were run using SPSS (IBM, Chicago, IL, USA) or Jamovi software (retrieved from https://www.jamovi.org) and graphed using Prism (GraphPad Software, San Diego, CA). Difference scores were calculated for the elevated plus maze (time spent in closed arms - time spent in open arms) and open field (time spent in periphery - time spent in center), and one-way analysis of variance (ANOVA) was used to determine whether difference scores varied by experimental condition. Tukey’s honestly significant difference (HSD) post hoc tests were used to examine significant omnibus tests for group differences. For the marble burying task, a two-tailed unpaired t-test was used to determine if the total number of buried marbles varied by hormonal condition. Two-way ANOVAs were used to determine if sucrose consumption and preference varied by hormonal condition.

One-way ANOVAs or Unpaired t tests were used to determine whether FosB cell counts varied by experimental conditions. Tukey’s HSD post hoc tests were used to examine significant omnibus tests for group differences. Unpaired t tests were used to determine whether FosB protein expression measured in Western blots differed between experimental conditions.

## Results

### Behavioral Assays of Anxiety

#### Elevated Plus

In C57BL/6 mice (Experiment 1), one animal was eliminated from the analysis because it jumped from the apparatus prior to the completion of the test.

There was a significant effect of hormone condition on the duration of time spent in the closed arms minus the duration of time spent in the open arms of the elevated plus maze (F(2,20) = 15.02, p = 0.0001). Post-hoc tests revealed that subjects in the no hormone group did not differ from estrogen-withdrawn animals in that both spent more time in the closed than open arms of the apparatus (p = 0.5964). However, estrogen-sustained animals differed significantly from both no hormone (p = 0.0018) and estrogen-withdrawn animals (p = 0.0001). Sustained animals spent a similar amount of time in the open and closed arms, indicating a lower-anxiety phenotype than no hormone or withdrawn females. The total distance traveled (F(2,20) = 0.5989, p = 0.5590) and average velocity (F(2, 20) = 0.1277, p = 0.8808) did not vary by hormone condition (**Figure 2**).

**Figure 2:**
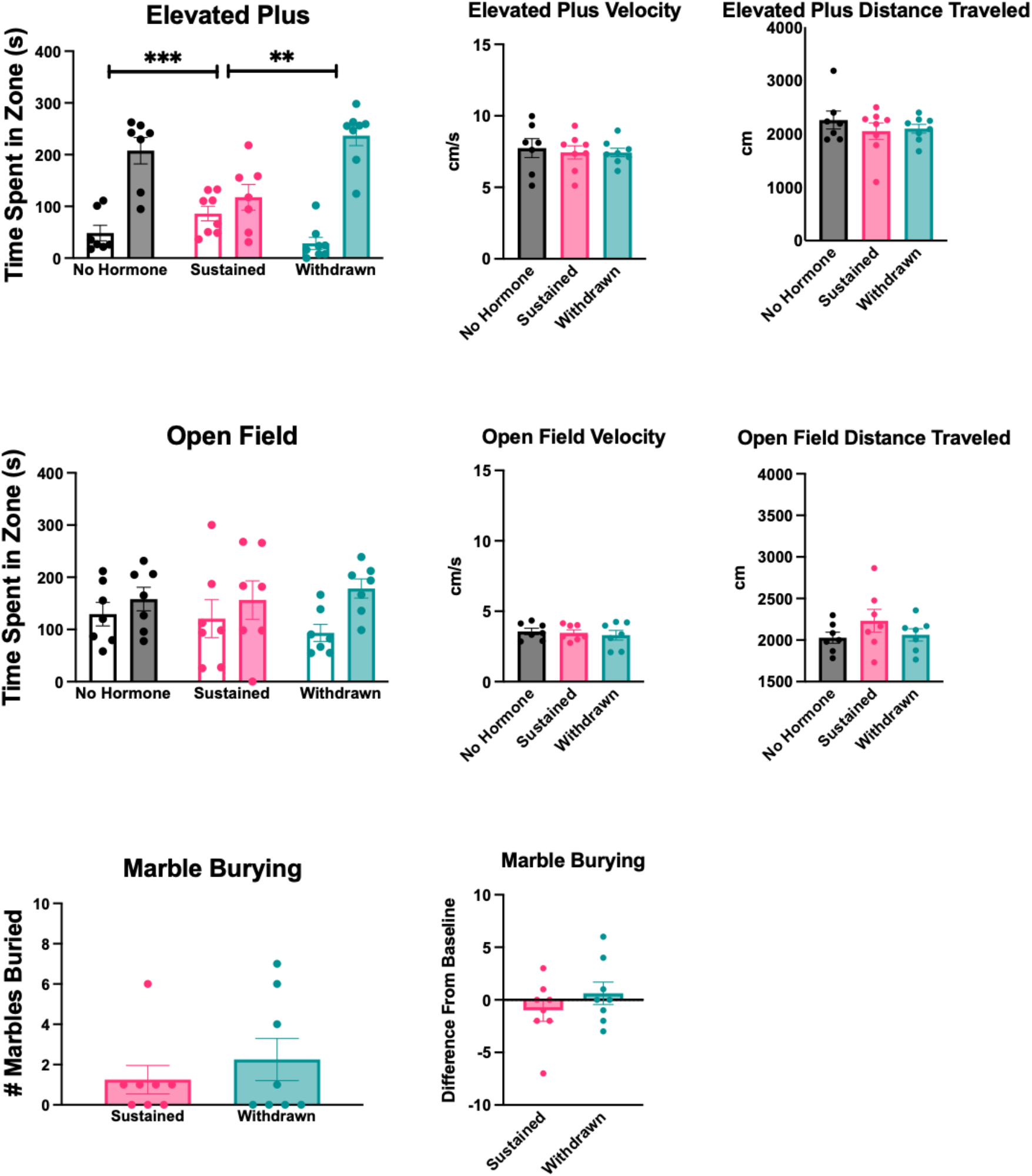
Impact of estrogen withdrawal on anxiety-like behaviors. In the elevated plus, estrogen-withdrawn and no hormone animals spend significantly more time in the closed arms of the apparatus (filled bars) than the open arms (open bars), indicating a higher-anxiety behavioral phenotype. In contrast, estrogen-sustained animals spent roughly equivalent amounts of time in the closed and open arms. In the open field, the time spent in the periphery (filled bars) vs. center (open bars) of the apparatus did not vary across hormone conditions. There was no effect of hormone condition on velocity or distance traveled in either the elevated plus or open field. Likewise, there was no significant difference between estrogen-withdrawn and estrogen-sustained animals in marble burying behavior. *** p < .0001, ** p < .001.

In transgenic reporter mice (Experiment 4), the duration of time spent in the closed arms minus the duration of time spent in the open arms was significantly higher in estrogen-withdrawn than estrogen-sustained females (t(14) = 3.476, p = 0.0037) (**Figure 5**). The total distance traveled (t(14) = 0.095, p = 0.926) and average velocity (t(14) = 0.090, p = 0.930) did not vary by hormone condition.

#### Open Field

In C57BL/6 mice (Experiment 1), there was no effect of hormone condition on the duration of time spent in the periphery minus the duration of time spent in the center of the open field (F(2,18) = 0.3488, p = 0.7102). The total distance traveled (F(2,18) = 1.227, p = 0.3166) and average velocity (F(2,18 = 0.2627, p = 0.7719) did not vary by hormone condition (**Figure 2**). In transgenic reporter mice (Experiment 4), there was no effect of hormone condition on the duration of time spent in the periphery minus the duration of time spent in the center of the open field (t(14) = 1.059, p = 0.3074) (**Figure 5**). The total distance traveled (t(14) = 0.547, p = 0.593) and average velocity (t(14) = 0.493, p = 0.630) did not vary by hormone condition.

#### Marble Burying

In C57BL/6 mice (Experiment 2),There was no effect of hormone condition on the number of marbles buried (t(14) = 0.7932, p = 0.4409). Likewise, there was no effect of hormone condition on the difference from baseline number of marbles buried (t(14) = 1.093, p = 0.2930) (**Figure 2**).

### Behavioral Assays of Motivation

#### Sucrose Preference

In C57BL/6 mice (Experiment 3), there was a significant interaction between hormone condition and test day on sucrose consumption (F(10,100) = 3.617, p < 0.001). During the acclimation period (Days 21-23), estradiol treatment increased sucrose consumption, reaching significance for the withdrawn group compared to the no hormone group on Day 21 (p = 0.003), both the sustained (p = 0.002) and withdrawn (p < 0.001) groups compared to the no hormone group on Day 22, and the sustained group compared the no hormone group on Day 23 (p = 0.030). During the testing period (Days 24-26), there was generally no effect of hormone treatment on sucrose consumption, although the estrogen-withdrawn group did consume more sucrose than the no hormone group on Day 24 (p = 0.031) (**Figure 3**).

**Figure 3:**
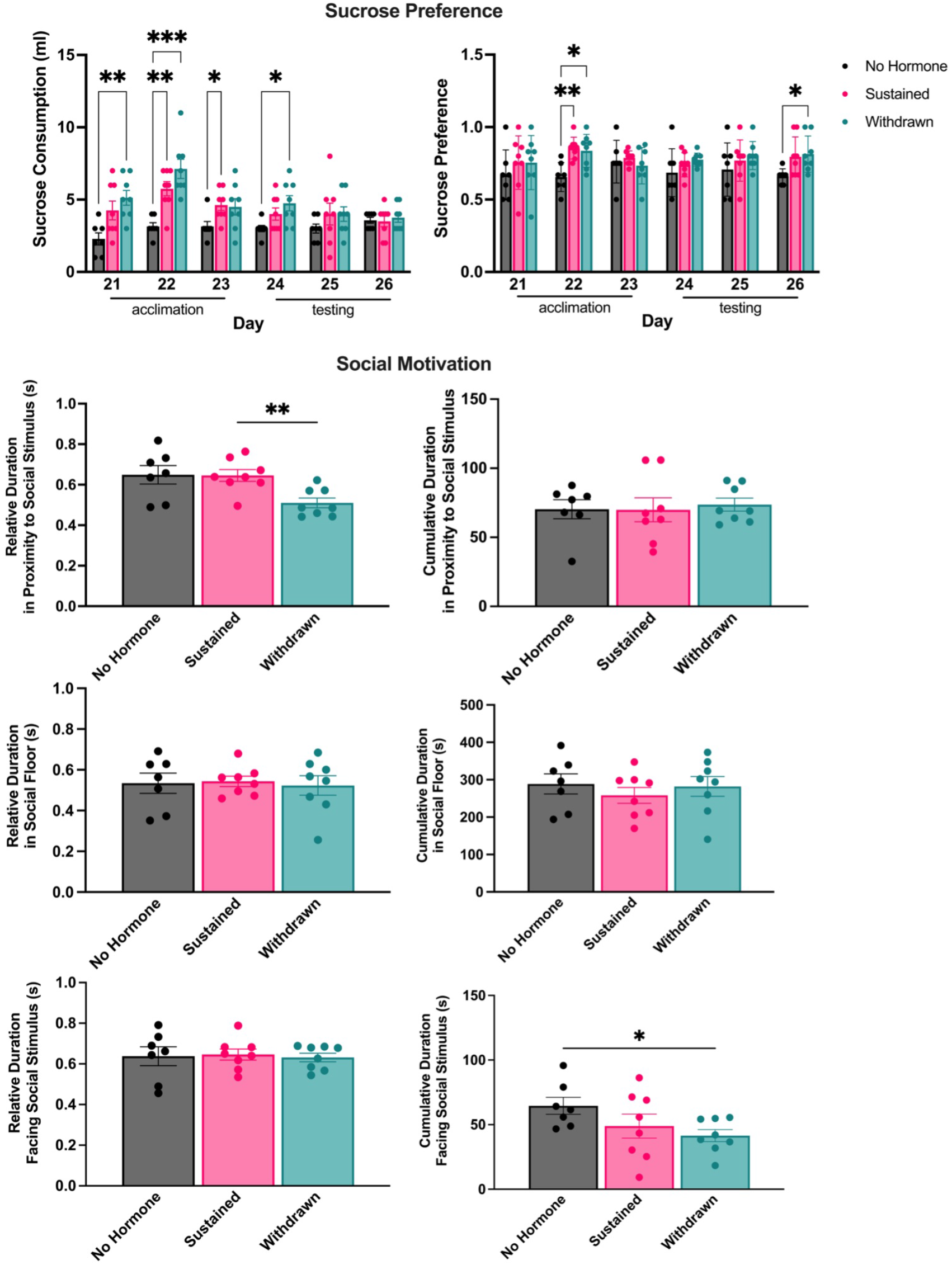
Impact of estrogen withdrawal on motivated behaviors. During the acclimation period, hormone treatment generally increased sucrose consumption in both sustained and withdrawn animals compared to no hormone females. This difference was maintained on the first day of the testing period, but sucrose consumption subsequently declined for both withdrawn and sustained animals. There was no consistent effect of hormone condition on sucrose preference, although estradiol treatment increased sucrose preference in both sustained and withdrawn animals compared to no hormone females on Day 22, and estrogen-withdrawn animals compared to the no hormone group on Day 26 of the testing period. In the social motivation test, estrogen-withdrawn animals spent less relative time in proximity to the social stimulus compared to sustained or no hormone animals, and less cumulative time facing the social stimulus than no hormone animals. Cumulative duration in proximity to a social stimulus, relative and cumulative time spent in the social floor, and relative time spent facing the social stimulus did not vary by hormone condition. *** p < .001, ** p < .005, * p < 0.05.

**Figure 4:**
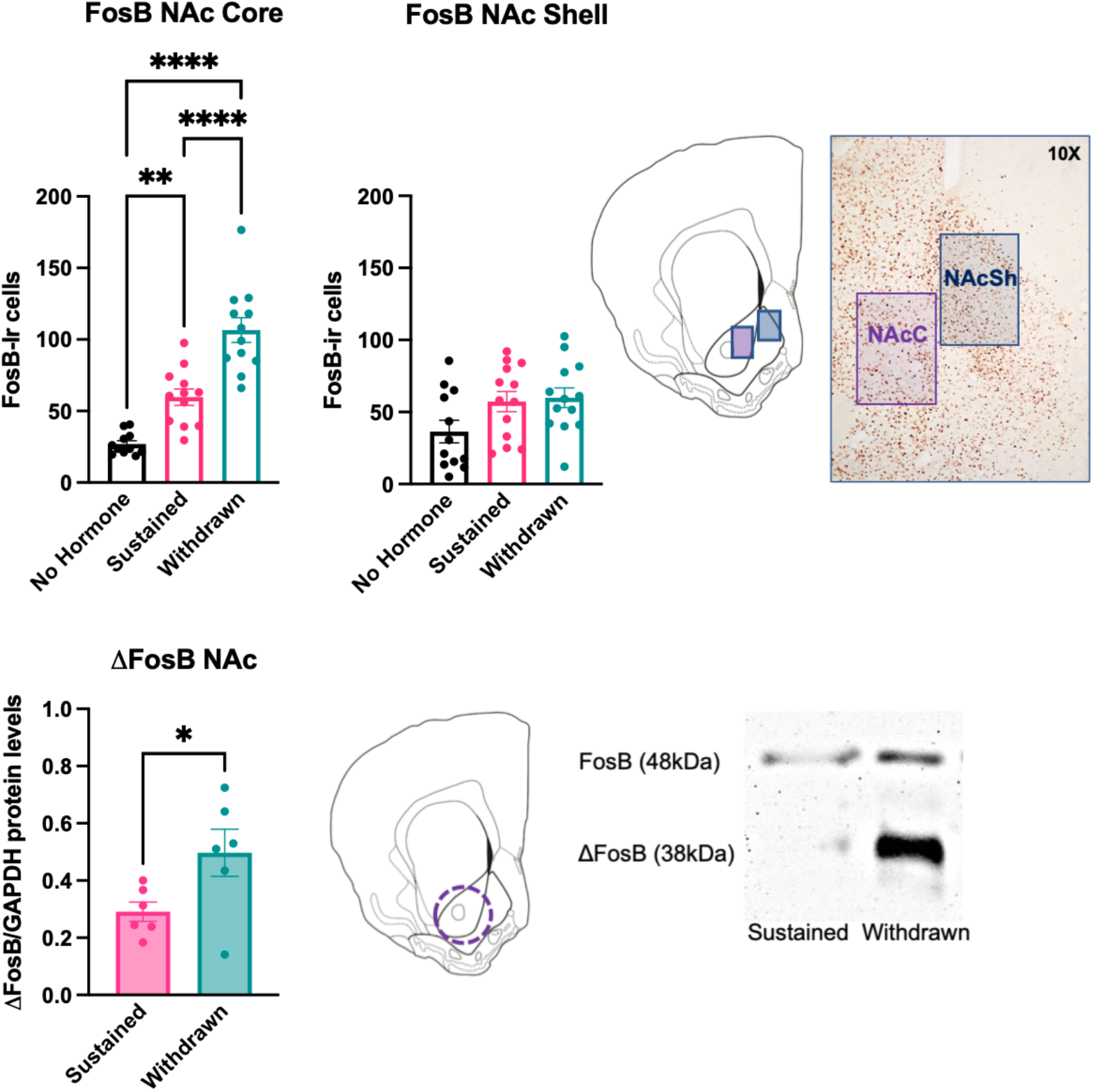
Estrogen Withdrawal increases ΔFosB in the NAc. On Day 28, animals in experiment 1 were euthanized via intracardial perfusion and their brains were processed for immunohistochemical localization of FosB protein in the NAc Core (NAcC) and (NAcSh). Estrogen-withdrawn females had significantly more FosB-immunoreactive (-ir) cells than both estrogen-sustained and no hormone animals in the NAc Core (NAcC). Estrogen-sustained animals also had more FosB-ir cells in the NAC than no hormone animals. There was no difference in FosB-ir cells across hormone conditions in the NAc Shell (NAcC). Cell counting domains and representative immunostaining shown to the right. In a subset of withdrawn and sustained animals from Experiment 1, brain tissue was processed for Western blots targeting ΔFosB in the NAc. Estrogen-withdrawn animals had significantly higher levels of ΔFosB protein than estrogen sustained animals. Representative core-biased tissue punch target and Western blot shown to the right. **** p < .0001, ** p < .001, * p < 0.05.

**Figure 5.**
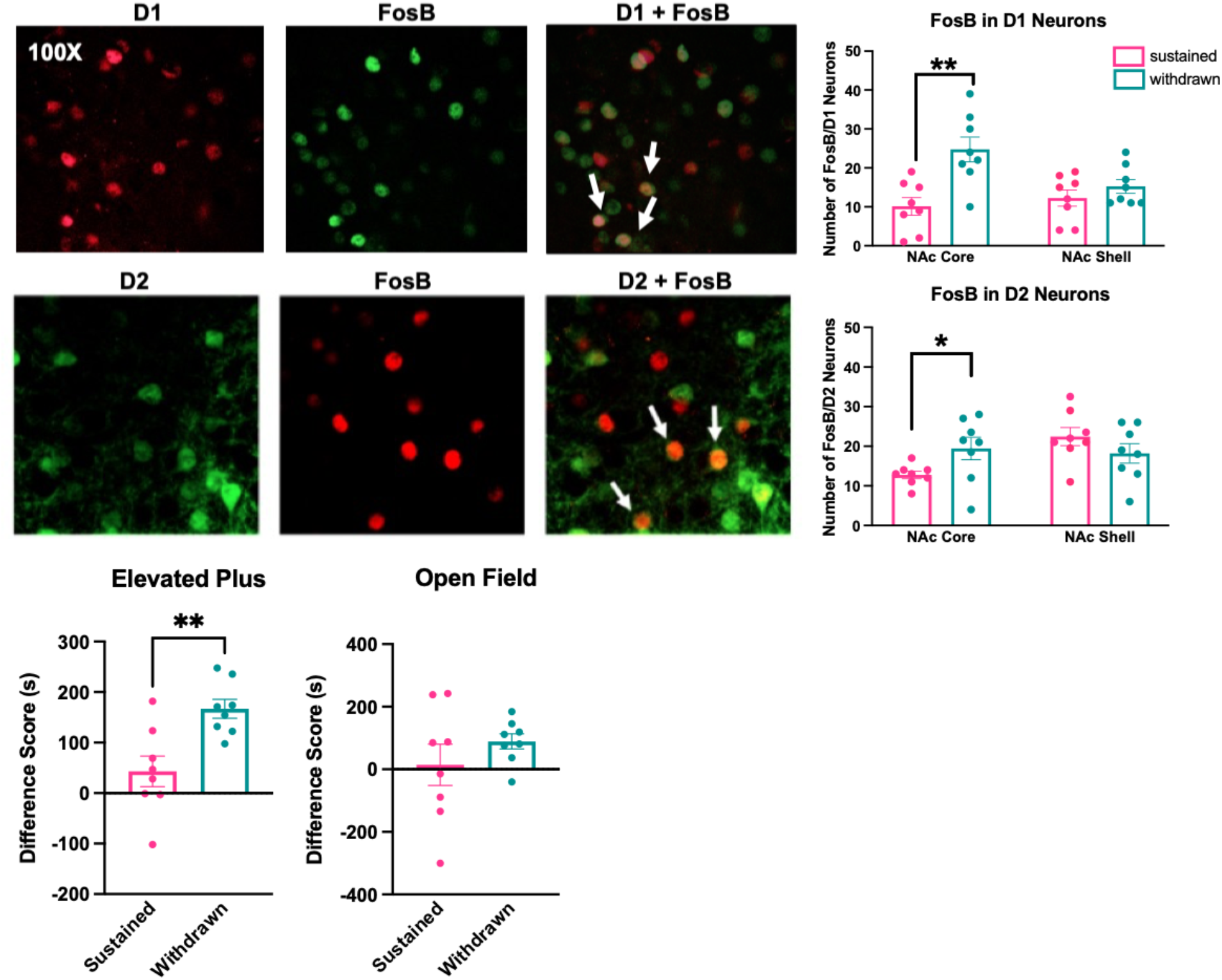
Estrogen withdrawal increases FosB-ir cells in D1 and D2 cells in the NAc core. In Experiment 4, *D1-tdtomato and D2-GFP* transgenic fluorescent reporter mice were used to examine whether FosB expression increased in a cell subtype-specific manner in the NAc. In *D1-tdtomato* mice, the number of double-immunofluorescent D1+FosB cells was increased in withdrawn animals compared to sustained animals in the NAcC, but not in the NAcSh. Similarly, tissue from *D2-GFP* mice showed that the number of double-immunofluorescent D2+FosB cells was increased in withdrawn animals compared to sustained animals in the NAcC, but not in the NAcSh. Representative single- and double-immunofluorescence shown above. Behavior in the elevated plus and open field was also assessed in transgenic reporter mice. Like in C57BL/6 mice, the duration of time spent in the closed arms minus the duration of time spent in the open arms of the elevated plus was significantly higher in estrogen-withdrawn than estrogen-sustained females. Also like in C57BL/6 mice, there was no effect of hormone condition on the duration of time spent in the periphery minus the duration of time spent in the center of the open field. ** p < .005, * p < 0.05.

There was a main effect of hormone condition on sucrose preference (F(2,20 = 5.827, p = 0.010). During the acclimation period (Days 21-23), estradiol treatment increased sucrose preference in both sustained (p = 0.003) and withdrawn (p = 0.016) animals compared to no hormone animals. During the testing period (Days 24-26), there was generally no effect of hormone treatment on sucrose preference, although estrogen-withdrawn group did have a significantly higher sucrose preference than the no hormone group on Day 26 (p = 0.027) (**Figure 3**).

#### Social Motivation

In C57BL/6 mice (Experiment 3), there was a significant effect of hormone condition on the relative duration of time spent in proximity to the social stimulus (F(2,20) = 5.835, p = 0.0101). Post-hoc tests revealed that estrogen-withdrawn animals spent significantly less time in proximity to the social stimulus than sustained (p = 0.009) or no hormone animals, although the later comparison did not reach statistical significance (p = 0.069). There was no effect of hormone condition on the relative duration of time spent in the social stimulus chamber (F(2,20) = 0.06187, p = 0.9392) or the relative duration facing the social stimulus (F(2,20 = 0.05164, p = 0.9498) (**Figure 3**).

There was no effect of hormone condition on the cumulative duration in proximity to the social stimulus (F(2,20) = 0.08677, p = 0.9172) or the cumulative duration of time spent in the social stimulus chamber (F(2, 20) = 0.4156, p = 0.6655). However, there was a significant effect of hormone condition on the cumulative duration facing the social stimulus (F(2,20) = 3.887, p = 0.049). Post-hoc tests revealed that estrogen-withdrawn animals spent significantly less time facing the social stimulus animal than no hormone animals (p = 0.044) (**Figure 3**).

### ΔFosB expression in the brain

#### Immunoreactivity in the NAc

Tissue from C57BL/6 mice showed that In the NAcC, there was a significant effect of hormone condition on the number of FosB-ir cells (F(2, 32) = 39.71, p < 0.0001). Post-hoc analyses revealed that both sustained (p = 0.0027) and withdrawn (p < 0.0001) animals had significantly more Fos-ir cells in the NAcC than no hormone animals. In addition, withdrawn animals had significantly more FosB-ir cells in the NAcC than sustained animals (p < 0.0001). In the NAcSh, however, there was no effect of hormone condition on FosB-ir cells (F(2,35) = 3.075, p = 0.8153 (**Figure 4**).

#### Western Blots in the NAc

Since FosB immunohistochemistry cannot distinguish between the two products of the FosB gene, ΔFosB and full-length FosB, Western blots were used to selectively measure ΔFosB protein levels in core-biased tissue punches from the NAc of sustained vs. withdrawn C57BL/6 females. Estrogen-withdrawn females had significantly higher ΔFosB levels (normalized to GAPDH) than did estrogen-sustained animals (t(10) = 2.309, p = 0.0436) (**Figure 4**). In contrast, there was no difference in levels of full-length FosB (t(10) = 0.2253, p = 0.8263).

#### Double-Immunofluorescence in D1 and D2 neurons in the NAc

Tissue from transgenic reporter mice showed that estrogen-withdrawal increased FosB-ir in both D1 and D2 neurons the NAcC, but not NAcSh. Tissue from *D1-tdtomato* mice showed that the number of double-immunofluorescent (FosB-ir + RFP-ir) cells was increased in withdrawn animals compared to sustained animals in the NAcC (t(14) = 3.723 p = 0.0011), but not in the NAcSh (t(14) = 1.097, p = 0.1456) (**Figure 5**). Similarly, tissue from *D2-GFP* mice showed that the number of double-immunofluorescent cells (FosB-ir + GFP-ir) was increased in withdrawn animals compared to sustained animals in the NAcC (t(14) = 2.248, p = 0.0206), but not in the NAcSh (t(14) = 1.273, p = 0.1119) (**Figure 5**).

## Discussion

Here we demonstrate for the first time that in C57BL/6 mice, estrogen withdrawal following HSP alters affective behaviors in the elevated plus, sucrose preference, and social motivation tests, but not in the open field or marble burying tests. These behavioral changes are consistent with increases in anxiety-like behavior and decreases in motivation that have been demonstrated in previous studies using HSP to model postpartum estrogen withdrawal, and extend the HSP model to the common C57BL/6 strain for the first time, providing valuable information about species and strain similarities and differences. In addition, we find that estrogen withdrawal following HSP increases ΔFosB in the NAc core, but not in the NAc shell. This increase occurs in both D1 and D2 receptor-bearing medium spiny neurons (MSNs). This novel finding suggests that ΔFosB may mediate long-term neuroplasticity in the NAc following postpartum estrogen withdrawal, and that these neuroplastic changes may be related to behavioral changes during this period.

Our finding that estrogen withdrawal following HSP produces a high-anxiety behavioral phenotype in the elevated plus is consistent with previous findings in ICR mice (Zhang et al. 2016; Yang et al. 2017), Sprague Dawley rats (Suda et al. 2008), and Syrian hamsters (Hedges et al. 2021). Findings in the open field following HSP are more variable both within (Yang et al. 2017; Zhang et al. 2016; Galea, Wide, and Barr 2001) and between (Hedges et al. 2021) species, suggesting that the elevated plus may be a more reliable thigmotaxis-based assay of anxiety-like behavior following HSP. A remaining question is whether estrogen treatment protects against anxiety or, in contrast, if estrogen withdrawal promotes anxiety? Our findings suggest that in C57BL/6 mice sustained high levels of estrogen may decrease anxiety, as estrogen-sustained females differed significantly from both no hormone, ovariectomized control females and estrogen-withdrawn females, whereas no hormone females did not differ from estrogen-withdrawn animals in the elevated plus.

We did not observe a significant effect of hormone condition on marble burying behavior, although marble burying was generally higher in estrogen-withdrawn animals than estrogen-sustained animals. Low levels of burying across all experimental groups may have made it difficult to detect differences. Nonetheless, the impact of peripartum hormone fluctuations on the compulsive subtype of anxiety bears further investigation given the increased risk of OCD in the peripartum period (Zambaldi et al. 2009) and the diagnostic challenges associated with peripartum OCD symptoms (Meltzer-Brody et al. 2018). Future experiments would benefit from using a different behavioral assay, such as the signal attenuation model of compulsive lever pressing. In this model, withdrawal from chronic administration of estradiol increases compulsive lever pressing (Flaisher-Grinberg et al. 2009), whereas blockade of D1 receptors decreases compulsive lever pressing (Joel 2006). These results, when coupled with the known role of the NAc in compulsive anxiety (Sturm et al. 2003), suggest that the signal attenuation model of compulsive lever pressing is a compelling model to test the role of postpartum estrogen withdrawal and NAc plasticity on compulsive anxiety behaviors.

In the current study, hormone treatment increased sucrose consumption during the acclimation phase when estradiol levels were high, but this effect declined for both estrogen-withdrawn and estrogen-sustained animals by the second and third days of the simulated postpartum period. This is consistent with previous findings in rats (Green, Barr, and Galea 2009), and may reflect a tolerance to continued sucrose presentation. In addition, mice in all hormone conditions reliably developed a sucrose preference, and hormone treatment significantly increased sucrose preference in both sustained and withdrawn females compared to no hormone females on the second day of the acclimation period. Although this preference was eliminated in both sustained and withdrawn animals on the first and second days of the simulated postpartum period, suggesting a possible anhedonic state, estrogen-withdrawn females again had a higher sucrose preference than no hormone females on the third day of the simulated postpartum period. These sucrose preference findings differ from previous findings in rats, in which only estrogen-withdrawn females exhibited a decrease in sucrose preference compared to females that continued to receive estradiol or an estrogen receptor beta (ERB) agonist (Green, Barr, and Galea 2009). However, our findings are consistent with recent data from Syrian hamsters, in which estrogen withdrawal following HSP did not impact sucrose preference (Hedges et al. 2021). Taken together, these data suggest that sucrose preference may be a species-dependent readout of motivation/anhedonia following HSP.

We also found that estrogen withdrawal following HSP decreased time spent in proximity to a social stimulus and facing a social stimulus, indicating a deficit in social motivation. This is the first time this has been established in an HSP model, but is consistent with a reduction in social motivation and other assays of behavioral despair in early postpartum female rats (Rincón-Cortés and Grace 2020). Our data suggest that deficits in social approach in naturally-parturient females may be mediated by estrogen withdrawal during the early postpartum period. Interestingly, in early postpartum female rats, deficits in social motivation and increased anxiety-like behavior in the elevated plus are correlated with decreases in the number of spontaneously active dopamine neurons in the ventral tegmental area (Rincón-Cortés and Grace 2020). This neural change is indicative of dopamine downregulation, or the deceased ability of the mesolimbic system to respond to behaviorally-relevant inputs. What’s more, it is consistent with microdialysis studies showing reduced baseline levels of extracellular dopamine in the NAc of early postpartum rats and ovariectomized females that are hormone-primed to induce maternal responsiveness (Afonso et al. 2009).

The induction of ΔFosB in NAc in response to chronic decreases in dopamine release is well-established in mice and humans. In mice, ΔFosB induction occurs selectively in D2 MSNs in response to chronic treatment with haloperidol, a D2 receptor antagonist (Lobo et al. 2013). In humans, ΔFosB induction occurs in NAc MSNs in response to dopamine depletion (Tekumalla et al. 2001), although the cell-type selectivity of this effect has not been established. Meanwhile, the induction of ΔFosB in D2 MSNs has been related to negative emotional states (Hamilton et al. 2018). These data map on to our current finding that estrogen-withdrawn females show increased ΔFosB expression in the NAcC, as measured by both Western blot and immunohistochemistry. We established that this increase in ΔFosB following estrogen withdrawal occurs in both D1 and D2 MSNs in the NAcC. Unlike the NAcSh, which is thought to be involved in shorter-term aspects of motivation, the NAcC is involved in the development of patterned motor programs to obtain rewards (Alcaro, Huber, and Panksepp 2007), such as might be expected following long-term physiological and/or neuroplastic changes. Interestingly, acute estradiol treatment decreases dendritic spine density in the NACc, but not NAcSh, in rats (Peterson, Mermelstein, and Meisel 2015) and Syrian hamsters (Staffend and Meisel 2012). The impact of chronic estradiol treatment and/or estrogen withdrawal on NAcC spine density is unknown, but the current results would predict an increase in dendritic spine density following postpartum estrogen withdrawal.

Evidence from functional neuroimaging studies in humans suggests that the striatum is dysregulated during postpartum psychiatric disorders. Specifically, people with postpartum depression have a blunted response in the striatum to positive, rewarding stimuli (Silverman et al. 2007), positive faces (Morgan et al. 2017), and their own infants’ joyful faces (Laurent and Ablow 2013). Despite this, very few studies have used animal models to investigate neuroplasticity in the mesolimbic pathway in relation to postpartum psychological states. In Flinders Sensitive Line (FSL) rats, a selectively-bred rodent model used to study depression, postpartum females show decreased extracellular dopamine levels in the NAc during pup interactions compared to Sprague-Dawley rats (Lavi-Avnon et al. 2008). Similarly, postpartum Sprague-Dawley rats that experienced chronic stress during gestation show reduced structural plasticity in the NAc, as well as reduced mobility in the forced swim test (Haim, Sherer, and Leuner 2014). However, this is the first study to examine the impact of estrogen withdrawal on molecular plasticity in the NAc mediating postpartum mood or affect. Future studies will benefit from this extension of the HSP model to C57BL/6 mice, in which genetic tools and behavioral assays can be leveraged to continue to investigate the role of postpartum estrogen withdrawal on mesolimbic plasticity and affective changes.

## Acknowledgements

Research supported in this publication was supported by the National Institute of Mental Health and the National Institutes of Health under Award Number R15MH125282 to LEB. The content is solely the responsibility of the authors and does not necessarily represent the official views of the National Institutes of Health. The authors wish to thank Ezekiel Mouzon (Icahn School of Medicine at Mount Sinai) for his assistance coordinating the shipping of transgenic reporter mice, Luke Troyon (Haverford College) for his assistance with confocal microscopy, and Victoria Lacorte and Victoria Bruce (Widener University) for their assistance in D1/FosB cell counts.

